# Two independent loss-of-function mutations in *anthocyanidin synthase* homeologous genes make sweet basil all green

**DOI:** 10.1101/2022.05.16.492104

**Authors:** Itay Gonda, Mohamad Abu-Abied, Chen Adler, Renana Milavsky, Ofir Tal, Rachel Davidovich-Rikanati, Adi Faigenboim, Tali Kahane-Achinoam, Alona Shachter, David Chaimovitsh, Nativ Dudai

## Abstract

Sweet basil, *Ocimum basilicum* L., is an important culinary herb grown worldwide. Although basil is green, many landraces, breeding lines and exotic cultivars have purple stems and flowers. This anthocyanins pigmentation is unacceptable in the traditional Italian basil. We used the recently published sweet basil genome to map quantitative trait loci (QTL) for flower and stem color in a bi-parental F_2_ population. It was found that the pigmentation is governed by a single QTL, harboring an anthocyanidin synthase (ANS) gene. Further analysis revealed that the basil genome harbors two homeologous ANS genes, each carrying a loss-of-function mutation. *ObANS1* carries a 1-bp insertion, and *ObANS2* carries a missense mutation within the active site. In the purple-flower parent, ANS1 is functional and ANS2 carries a nonsense mutation. The functionality of the active allele was validated by complementation in an Arabidopsis ANS mutant. Moreover, we have restored the functionality of the missense-mutated ANS2 using site-directed activation. We found that the non-functional alleles were expressed to similar levels as the functional allele, suggesting polyploids invest futile effort in expressing non-functional genes, harming their superior redundancy. We show here we can harness basil’s genomics and genetics to understand the basic mechanism of metabolic traits.

## Introduction

Basil, *Ocimum basilicum* L., of the *Lamiaceae* family, is a leading aromatic crop in agricultural fields and home gardens. It belongs to the genus *Ocimum*, which comprises up to 160 different species (Paton *et al*., 1999). *O. basilicum* harbors a large diversity among its genotypes depicting unique aromas, leaf sizes and shapes, leaf and stem color, inflorescence color and structure, grew habit and seeds morphology (Dudai and Belanger, 2016). These diverse genotypes have a wide range of uses, primarily as a culinary herb and as a source for essential oils and ornamentals. ‘Genovese’ basil, the type of basil used in the Italian Pesto sauce, is the most widespread basil grown globally. While commercial ‘Genovese’ basils are all green, markets also display purple basils or basils with purple stems and flowers.

Anthocyanins are a group of water-soluble pigments conferring purple, red and blue colors in multiple plants (Holton and Cornish, 1995). They exist in almost all plant species except from *Caryophyllales* that accumulate betalains pigments instead (Polturak and Aharoni, 2018; Tanaka *et al*., 2008). The biosynthesis of anthocyanins is a multistep pathway that starts from the amino acid L-phenylalanine (Fig. 1). Several colorless or yellow intermediates precede the anthocyanins. The first enzyme that generates purple/blue/red pigments is anthocyanidin synthase (ANS; synonym: leucoanthocyanidin dioxygenase, LDOX), which oxygenates the leucoanthocyanidin substrate and generates colored anthocyanidin aglycons (Falcone Ferreyra *et al*., 2012). Next, multiple sugar substitutions on various residues produce various colorful anthocyanins.

**Fig. 1.**
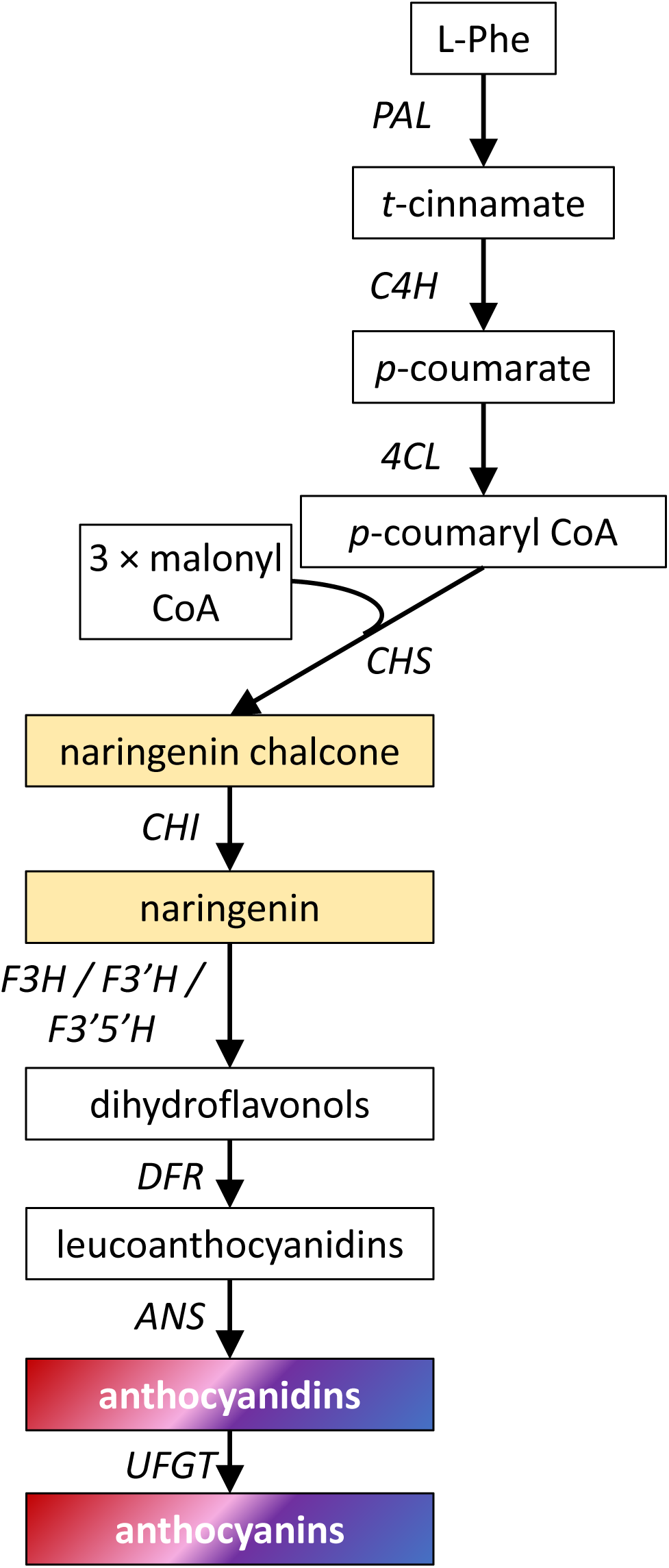
Biosynthesis pathway of anthocyanins. PAL, phenylalanine ammonia lyase; 4CL, p-coumarate CoA ligase; C4H, coumarate 4-hydroxylase; CHS, chalcone synthase; CHI, chalcone isomerase; F3H, flavanone 3-hydroxylase; F3′H, flavonoid 3′-hydroxylase; F3′5′H, flavonoid 3′,5′-hydroxylase; DFR, dihydroflavonol 4-reductase; ANS, anthocyanidin synthase; UFGT, UDP-glucose flavonoid 3-O-glucosyltransferase. Enzyme names are in italic letters.

In basil, purple pigmentation is a common feature in the leaves, flowers and stems (Carović-Stanko *et al*., 2011). While the term purple basil usually refers to the color of the leaves, green-leaf basils with purple flowers and stems are common (Phippen and Simon, 1998). Various anthocyanins were characterized from basil plants, with cultivar being a major factor determining their levels (Flanigan and Niemeyer, 2014; Phippen and Simon, 1998; Prinsi *et al*., 2019). The accumulation of anthocyanins in basil was also highly dependent on plant age (McCance *et al*., 2016), peaking at flowering time (Phippen and Simon, 1998). Fall-grown purple basil (cv. Dark Opal) accumulated significantly higher levels of anthocyanins than summer-grown (Nguyen and Niemeyer, 2008). Phippen and Simon (2000) showed that two dominant alleles govern the inheritance of basil color by using a complete diallel cross in segregating F_2_ individuals. They documented a high level of instability of purple leaf color seen in spotted or green/purple intermediate phenotypes of the offspring. That instability was also observed in vegetative cuttings of purple basils, where the position of the cutting influenced its color retaining (Phippen and Simon, 2000).

This project aimed to understand the molecular basis of anthocyanins accumulation in sweet basil. We used a previously-developed F_2_ population derived from a cross between a green cultivar and a purple-flower cultivar. Using genotyping-by-sequencing, we showed that a single quantitative trait loci (QTL) governs the color tait. Two mutations in the two ANS homelogous genes were validated. Finally, we raise the question of whether polyploids pay penalties when they express more non-functional genes than diploids, eliminating their hypothesized advantage of redundancy.

## Materials and Methods

### Plant material

The F_2_ mapping population was grown in greenhouse conditions as described in Gonda *et al*. (2022). Flower and stem color were visually evaluated on each F_2_ plant in the greenhouse on the emergence of the first flower. For gene expression analysis, 3 plants of ‘Perrie’ cultivar and 3 plants of ‘Cardinal’ culivar were grown in open field conditions with drip irrigation. Leaves samples were taken at 10 leaf-pairs stage, from the 7^th^ and 8^th^ pairs. Flowers were sampled after 2 months.

### DNA extraction, genotyping by sequencing and association mapping

DNA extraction, GBS libraries construction, SNP calling and association mapping were described in Gonda *et al*. (2022).

### Linkage groups determination

Linkage groups (LGs) were built with JoinMap v4.1. Briefly, of the 23,411 SNPs detected, only sites where both parents were homozygous continued the linkage analysis (using Tassel). The data was further filtered with JoinMap, and sites that did not show a disomic distribution of 1:2:1 were filtered out, and only one locus was kept when two loci were > 95% similar. Then, when adjacent sites on the same scaffold were distant less than 2.13 Mbp (on average), only the site with less missing data was kept. Linkage analysis was performed with JoinMap using regression mapping with the Cosambi map function. The parameter used for grouping was recombination frequency from 0.5 to 0.05 with a step of -0.05. LGs were set, and map distances were calculated based on the grouping tree with the regression mapping function. Homeologous LGs were determined considering the BUSCO analysis of Gonda *et al*. (2020) that defined homeologous scaffolds.

### RNA-sequencing

RNA sequencing, including RNA extraction, library preparation and gene expression analysis was performed as described in Gonda *et al*. (2020). In addition to the ‘Perrie’ cultivar, we also used the ‘Cardinal’ cultivar.

### Sequencing *ObANS* genes

Flowers RNA was converted to cDNA using a synthesis kit (PCR Biosystems, https://pcrbio.com/usa/) after DNAse treatment (Thermos Fisher scientific, https://www.thermofisher.com/il/en/home.html). Due to high identity between the homeologous copies, the ANS genes were amplified using a single pair of primers: F 5’ ATGGTTGCTTCAATTACGGCA 3’; R 5’ CAACTAGATTTATCATCAACCACCACC 3’. The PCR products were cloned into pJET vector (Thermos Fisher scientific) and transformed into DH5α competent cells plated on LB medium containing ampicillin for selection. Ten individual colonies were grown overnight, the plasmids were extracted, and the inserts were sequenced from all colonies to address all possible insert fragments. To restore the functionality of the H292Q mutation of *ObANS1_Perrie* we used the following primers: F 5’ CAAAAGCATTCTGCACCGCGCCTCCGTCAA 3’; R 5’ TTGACGAGGCCGCGGTGCAGAATGCTTTTG 3’.

### Complementation test of *ObANS* genes in Arabidopsis

All *ObANS* genes were inserted into the pBI121 plasmid under the control of the constitutive Cauliflower mosaic virus 35S promoter using the SacI and XbaI restriction enzymes (Thermo-scientific) and the NEBuilder ligation system (NEB). The plasmids were transformed into *Agrobacterium tumefaciens* strain GV3101 using electrotransformation and positive colonies were selected with kanamycin. Agrotransformation of Arabidopsis plants with T-DNA insertion at the *AtANS* gene (SALK_073183, Ohio State University Arabidopsis Biological Resource Center, https://abrc.osu.edu/), were performed using the floral dip technique (Clough and Bent, 1998). The F1 seeds then germinated on a kanamycin-containing MS medium, and seedlings harboring the plasmid were grown, followed by F_2_ seeds collection. The F_2_ plants were grown on a kanamycin-containing MS medium at 20°C with a 12h photoperiod.

### Anthocyanins extraction

Flowers tissues of 3 plants of both ‘Perrie’ and ‘Cardinal’ cultivars were collected and flash-frozen in liquid N_2_. Samples were then ground to uniform powder. Afterward, 100 mg of tissue was weighed, 200 μl of 80% methanol was added, and samples were vigorously vortexed. Samples were then sonicated for 20 min at RT followed by 10 min centrifugation at 21,000 *g* at 4°C. The extraction procedure was repeated twice. Finally, the samples were filtered through a syringe filter of 0.22 μm (GHP Membrane, PALL, USA) to new amber vials.

### Anthocyanins analysis

Liquid chromatography/time of-flight/mass spectrometry (LC-TOF-MS) analysis was carried out on an Agilent 1290 Infinity series liquid chromatograph coupled with an Agilent 1290 Infinity DAD and Agilent 6530C Accurate Mass quadrupole Time of Flight (qTOF) mass spectrometer (MS) (Agilent Technologies, Santa Clara, USA). Compounds were separated on a Zorbax Extend-C18 Rapid Resolution HT column (2.1 × 50 mm, 1.8 μm; Agilent Technologies). The gradient elution mobile phase consisted of H_2_O with 0.1% (v/v) formic acid (eluent A) and acetonitrile containing 0.1% (v/v) formic acid (eluent B). The column was equilibrated with 1% B at a flow rate of 0.3 mL × min^-1^ for 1 min, then increased to 80% B by the following steps: 2-3 min, 20% B; 4-5 min, 30% B; 10-11 min, 50% B; 12-13 min, 80% B. Column was washed with 95 % B at 14 min for 1 min and readjusted to 1% B for 2 minutes. The eluted compounds were subjected to Jet Stream electrospray ionization interface (ESI) operated in positive mode with the following settings: 8 L × min^-1^ gas at 300°C, 35 p.s.i. nebulizer pressure, 10 L × min^-1^ sheath gas at 300°C, capillary voltage (VCap) of 3,000 V, fragmentor to 140 V, and skimmer to 65 V. Data was collected from mass/charge (m/z) ratio of 100-1,700. The flow rate of the mobile phase was 0.3 mL × min^-1^ and the column oven temperature was 30 °C. The main (therefore representative) ions formed in ESI source (mainly [M]^+^; [M+H]^+^; [M+Na]^+^) of target compounds were detected using the ‘find compound by formula’ function and analyzed by Masshunter qualitative and quantitative analysis software version B.07.00 (Agilent technologies). For untargeted analysis, the platform of MPP (Mass profiler professional, Agilent Technologies) was used. In total, 432 compounds were integrated using molecular feature of which 107 showed > 2 fold change between ‘Perrie’ and ‘Cardinal’ samples (Moderate *t*-test, p < 0.05). Compounds were annotated using IDBrowser based on exact mass compared to METLIN Metabolite and Chemical Entity Database.

### ANS sequence analyses and model predictions

Sequence comparisons and alignments of both nucleotides and amino acids were done with Clustal Omega (https://www.ebi.ac.uk/Tools/msa/clustalo/) using default parameters and visualized by BoxShade 3.21 (https://embnet.vital-it.ch/software/BOX_form.html). Evolutionary conservation scores were calculated by Consurf (Ashkenazy *et al*., 2016) for each of the ANS variants. ANS models were predicted using RaptorX (Källberg *et al*., 2012). AtANS (PDB 1gp6) was used as a template (Wilmouth *et al*., 2002). For electrostatic surface potential, hydrogens were added using PROPKA (Olsson *et al*., 2011) followed by calculation of the electrostatic potentials by APBS (Dolinsky *et al*., 2007). Secondary structure fold prediction was done by HHpred (Söding *et al*., 2005).

## Results

### Phenotyping the population for color traits

Many basil landraces and accessions have pink/purple flowers, while most cultivars found on markets have white/green flowers. We used a bi-parental population derived from a cross between the Genovese cultivar ‘Perrie’ and the ornamental cultivar ‘Cardinal’ (Dudai *et al*., 2018) to study the genetic basis for color traits. ‘Perrie’ has a green stem, green bracts and sepals, and white petals, while ‘Cardinal’ has a purplish stem, deep purple bracts and sepals and pink petals (Supplementary Fig. S1 at *JXB* online). The purple color results from the presence of four different anthocyanins with various glycoside substitutions, as was observed by LC-TOF analysis (Supplementary Table S1 at *JXB* online). The F_1_ plants had purple flowers but not as intense as the ‘Cardinal’ flower (Supplementary Fig. S1). Next, 173 F_2_ segregants were grown and visually scored for the color of the flower parts and the stem. The flower and stem color segregated in the F_2_ offspring and displayed varying intensities of purple color. There was an agreement between all non-colored individuals in all traits; offspring with white petals had green stems, bracts and sepals. χ^2^ distribution analysis showed that the ratio between the white/green phenotype to the pink/purple phenotype fits a 3:1 single dominant gene inheritance model (Table 1). A look at the intensities of the purple color suggested an intermediate phenotype exists, subdividing the purple phenotype to deep- and light-purple (Table 1; Fig. 2). Although there was no complete agreement on purple intensities among sepals, stem and bracts, contingency analysis within purple segregants only, indicated that the color intensities among these tissues are related to each other (Supplementary Table S2). We have further tested the possibility that the intermediate purple phenotypes resulted from a single gene in an incomplete dominance model. The colors of the bracts, but not the other tissues, were segregated in a 1:2:1 ratio according to the χ^2^ probability test (Table 1).

**Table 1.**
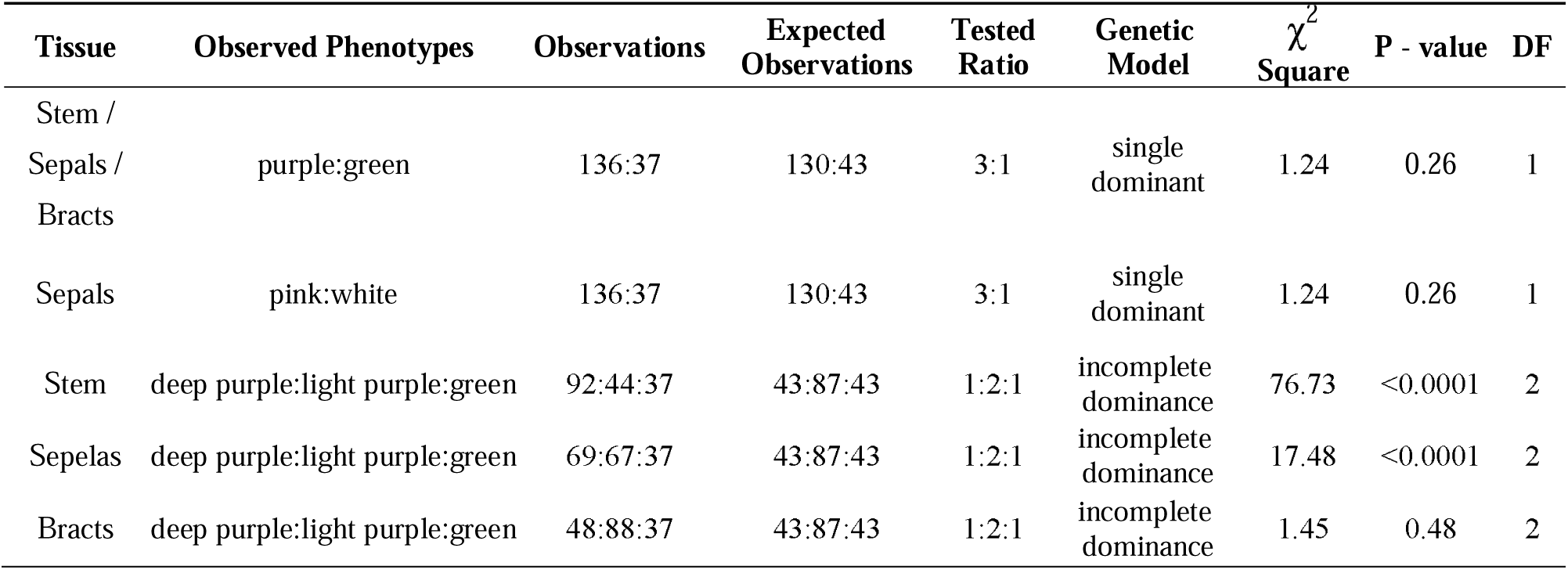
Observed and expected colors phenotype in the research population

**Figure 2.**
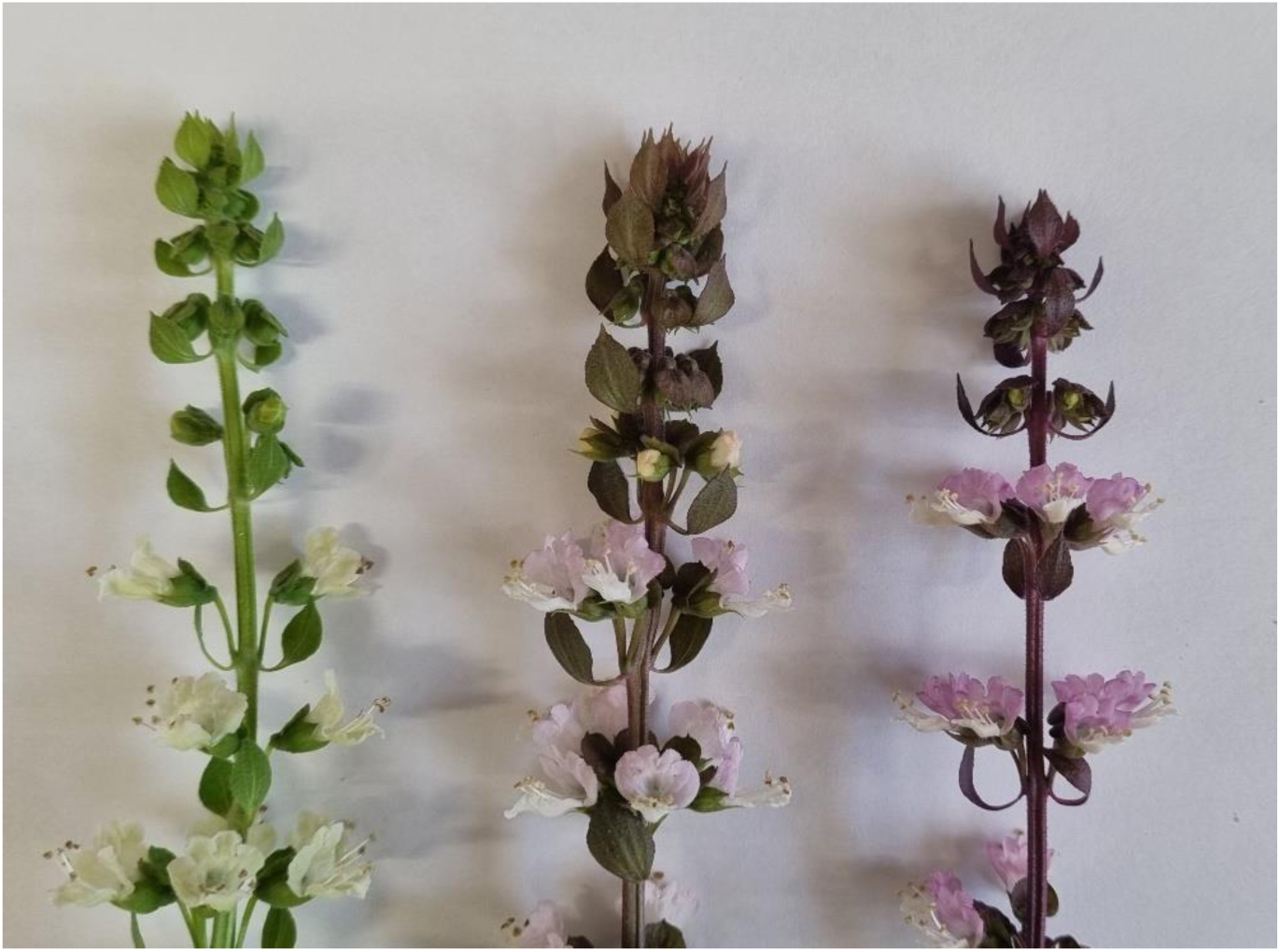
Color phenotype within the F2 population. Three representative color phenotypes of the F2 offspring: green with white sepals (left), light purple with pink petals (middle), deep purple with purple petals (right).

### Population’s genotyping and linkage groups construction

The population was genotyped with GBS, as described in Gonda *et al*. (2022). Briefly, 23,411 polymorphic sites between ‘Perrie’ and ‘Cardinal’ were generated for the mapping population. The heterozygosity levels were 9% for ‘Perrie’, 33% for ‘Cardinal’, and the population’s mean was 51% ± 0.7.

We next build linkage groups (LGs) and a genetic map based on the SNPs data. To reduce the complexity of the data, we have further filtered the SNPs to have a minimum genotyping depth of 10. The generated ABH genotype (A, ‘Perrie’; B, ‘Cardinal’) contained 11,857 sites (excluding sites where at least one of the parents was heterozygous). The data was further filtered with JoinMap, and sites that did not show a disomic distribution of 1:2:1 were filtered out, and only one locus was kept when two loci were > 95% similar. Then, when adjacent sites on the same scaffold were distant less than 2.13 Mbp (on average), only the site with less missing data was kept. This rigorous filtering resulted in 867 sites spreading over 152 scaffolds. We detected 24 linkage groups, but the average size was only 24 cM (SD = 21.3cM), and the median was 14 cM. Moreover, the SNPs did not order sequentially according to their scaffolds within most of the linkage groups (see examples in Supplementary Fig. S2). Hence, we only divided the scaffolds into LGs and did not continue to pseudomolecules scaffolding. Based on the BUSCO analysis performed by Gonda *et al*. (2020), we have determined homeology between LG couples that were arbitrarily classified into subgenomes A or B (Supplementary Table S3). We also detected chimeric scaffolds that span over 2 or 3 LGs (Supplementary Tables S3-4).

### Association mapping of the color traits

Due to the high heterozygosity level of the ‘Cardinal’ parent and the short genetic sizes of the linkage groups generated, we performed an association analysis rather than QTL mapping. To reduce complexity and false positives, we filtered the data to sites with DP > 15. That resulted in 8,496 polymorphic sites (including sites where one of the parents was heterozygous), which were checked for associations with the various color phenotypes. The results showed that no matter which color trait was tested, three scaffolds were strongly associated with the color traits: 393, 2608 and 7350 (Fig. 3). These scaffolds belong to the same LG (LG 4A), which suggests that the QTL at LG 4A spans over the entire chromosome. Moreover, also scaffold 120, which belongs to LG 4B, showed association with all color traits. We then checked whether the alleles at the QTL (position 7350_2,452,081) also contribute to the incomplete dominance we observed for bracts color phenotype (Fig. 4). Nonparametric comparisons of each allele-pair using Wilcoxon analysis indicated that the heterozygous status is significantly different from each homozygous status (*p*-value <0.0001). The phenotypic variance explained by this locus is 62%, according to contingency analysis test.

**Fig. 3.**
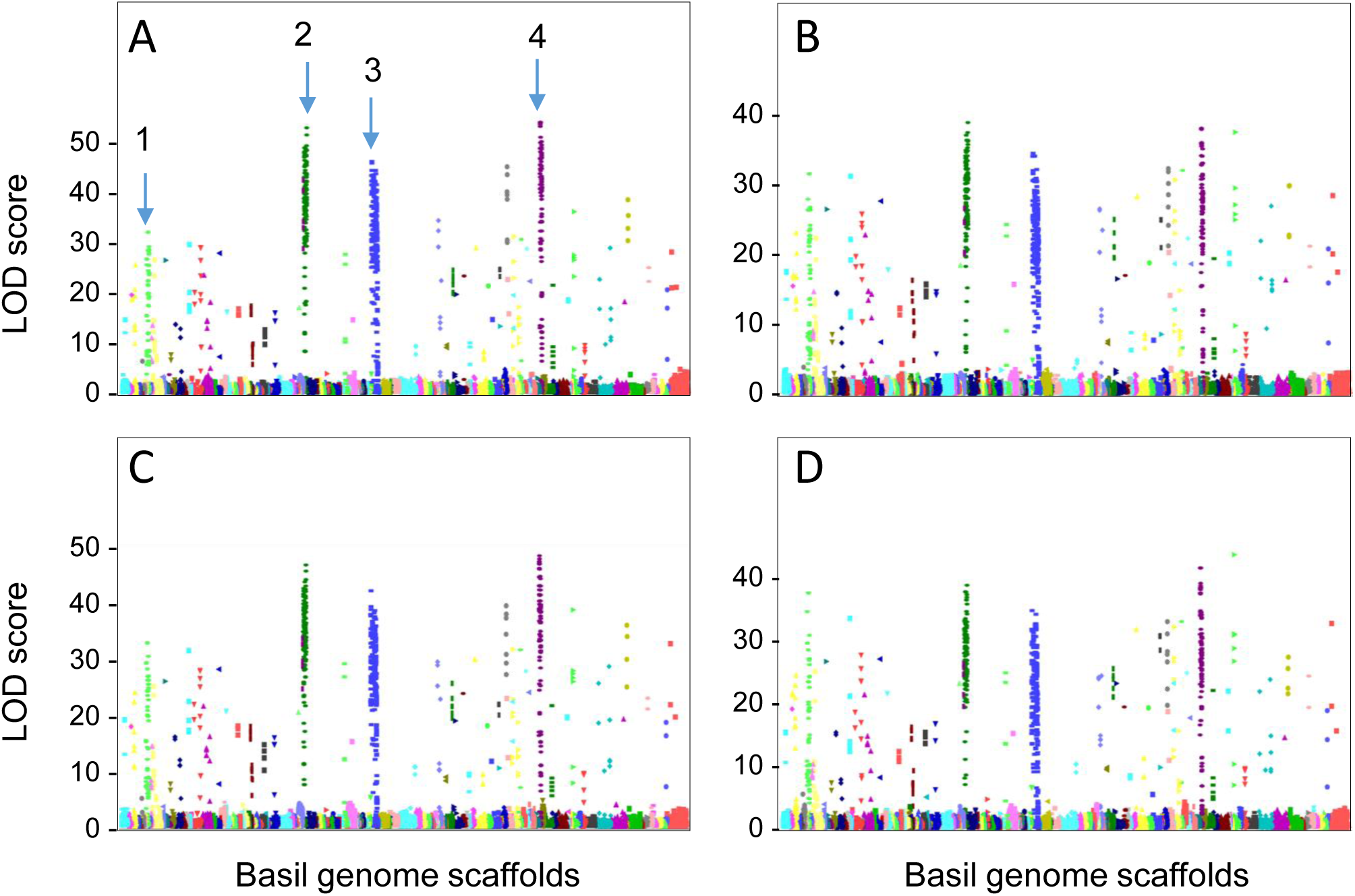
Association mapping of basil color traits across sweet basil genome scaffolds. Manhattan plots of the QTL for the color of: A. Bract leaves, B. Sepals, C. Stem, D. Petals. The associations of the traits and SNP within the F2 population were calculated using Tassel with GLM algorithm. Different colors and symbols represent the different scaffolds. Numbers represent: 1, scaffold 120; 2, scaffold 2608; scaffold 393; 4, scaffold 7350.

**Figure 4.**
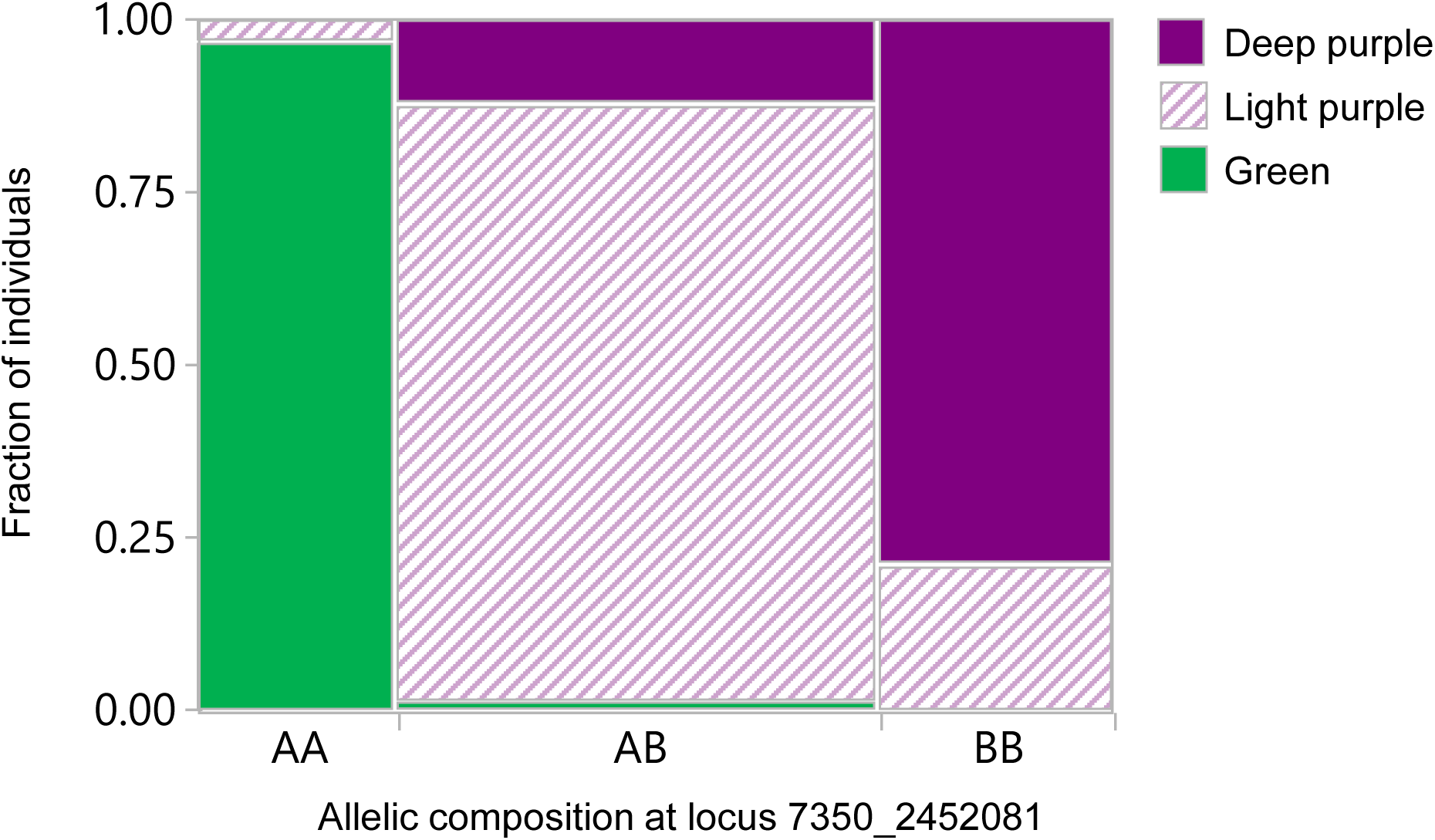
Contingency analysis of the bract leaves color phenotype vs. the allelic composition at site 7350_2452081. The width of each tile is proportional to the number of segregants with a given allelic composition in scaffold 7350 at position 2,452,081 bp. The height of each tile represents the fraction of the segregants with a given color with a given allelic composition. n=173.

### Resolving flower and stem color QTL

The observed intervals of several to dozens Mbp make it challenging to resolve the QTL and find the causative molecular factor using forward genetics only. To overcome that hurdle, we have adopted a reverse genetics approach with candidate genes from the anthocyanins biosynthetic pathway. We used tblastn algorithm to scan the basil genome for the entire anthocyanins biosynthetic pathway genes. We found that the entire pathway is duplicated in the ‘Perrie’ genome with copies in homeologous scaffolds (Supplementary Table S5). The only gene found in the color QTL scaffolds was the gene encoding for anthocyanidin synthase (ANS) enzyme. One copy of ANS gene was found in scaffold 7350 (termed *ObANS1*) and another copy was found in scaffold 120 (termed *ObANS2*) (Supplementary Table S5). Both genes were predicted to include two exons and are 96% identical in the nucleotide level, the intron however, is more divergent with 85% identities. Two molecular scenarios can explain the no anthocyanins phenotype of ‘Perrie’ flowers: 1) the expression of both ANS copies is suppressed in ‘Perrie’ due to *cis*-acting elements, plausibly located in the promoter; 2) ‘Perrie’ copies of ANS carry mutations in comparison to ‘Cardinal’ altering their activity. The expression levels of both *ObANS* genes was monitored in the flowers and the leaves of both genotypes using RNA-seq analysis. In both cultivars, both *ANS* genes were highly expressed in the flowers and hardly expressed in the leaves (Fig. 5). *ObANS2* was not differentially expressed between the two cultivars. *ObANS1* showed significantly higher expression levels in ‘Cardinal’ flowers than in ‘Perrie’ flowers. Yet, this difference in expression cannot explain the no anthocyanins phenotype of ‘Perrie’ flowers. Interestingly, both ‘Perrie’ and ‘Cardinal’ expressed the entire homeologous sets of anthocyanins biosynthetic genes (Table S6).

**Figure 5.**
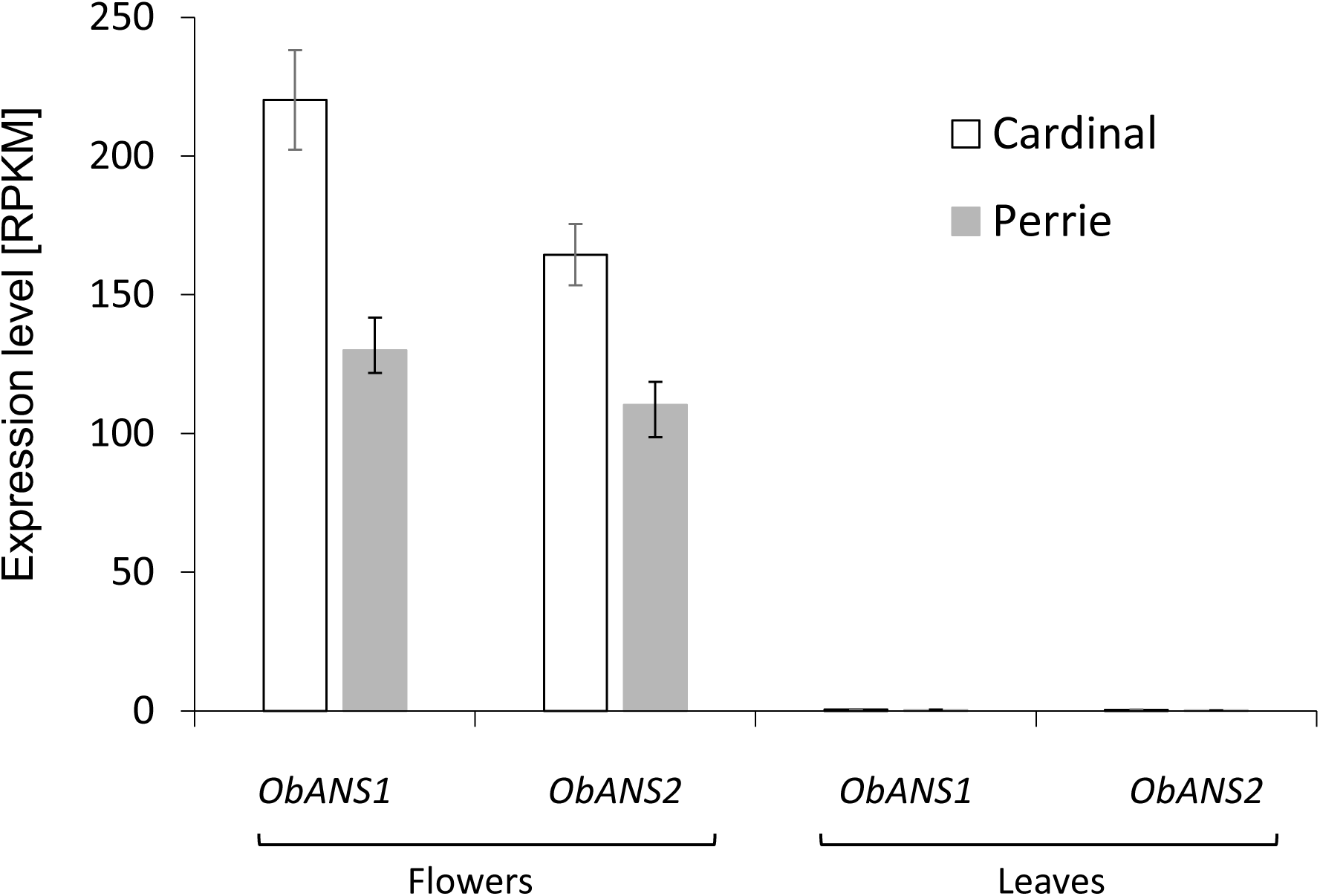
Expression levels of ANS genes in sweet basil. The expression of *ObANS1* and *ObANS2* in the parental lines, ‘Perrie’ and ‘Cardinal’, in the flowers and leaves was determined using RNA-seq. Values are mean of 3 biological repeats ± SEM.

### *ObANS* enzymes sequence analysis

Next, we looked for possible causative polymorphic sites between ‘Perrie’ and ‘Cardinal’. For that, we have extracted and sequenced the coding region of the ANS genes from cDNA of ‘Perrie’ and ‘Cardinal’ flowers. ANS1_Perrie carries a 1-bp deletion at position 993 bp (Supplementary Fig. S3, red shaded) in comparison to ANS1_Cardinal and ANS2 from both cultivars. ANS1_Cardinal carries a 9-bp deletion after 1,077 bp (Supplementary Fig. S3, light blue shaded). Several other SNPs were evident among the four ANS genes with 96.3 to 98.1 % identities (Supplementary Table S7). To understand the functional outcome of these deletions and SNPs, we have aligned basil ANS protein sequences along with the protein sequence of the Arabidopsis ANS (AtANS), that its’ structure was resolved using X-ray crystallography (Wilmouth *et al*., 2002). The 9-bp deletion in ObANS1_Cardinal did not seem to cause deletion in functionally important amino acids. All the critical amino acids according to AtANS were conserved in ObANS1_Cardinal (Fig. 6), suggesting it encodes a functional enzyme. In ANS1_Perrie, the 1-bp deletion did not cause a premature stop codon, yet the frameshift causes major variations in the last 58 amino acids of the protein (Fig. 6). That resulted in a protein with only ∼83% amino acid identities with other basil ANSs that are 95-98% identical (Supplementary Table S8). According to the crystal structure of AtANS, this arm of the enzyme harbors four amino acid residues that are important to the binding of the substrate. In ANS1_Perrie, these four amino acids were not conserved due to the frameshift plausibly altering its functionality. ObANS2_Cardinal carries a G ⟶ T mutation at position 460 (Supplementary Fig. S3, yellow shaded), resulting in a premature stop codon after only 153 amino acids that are 211 amino acids short of the full protein (Fig. 6). Finally, ObANS2_Perrie carries a C ⟶ G mutation at position 876 (Supplementary Fig. S3, green shaded), resulting in an amino acid substitution at position 292, replacing histidine with glutamine (Fig. 6). That H292Q mutation is at a histidine residue that is one of the three residues docking the Fe^++^ ion in the enzyme’s active site (Wilmouth *et al*., 2002).

**Figure 6.**
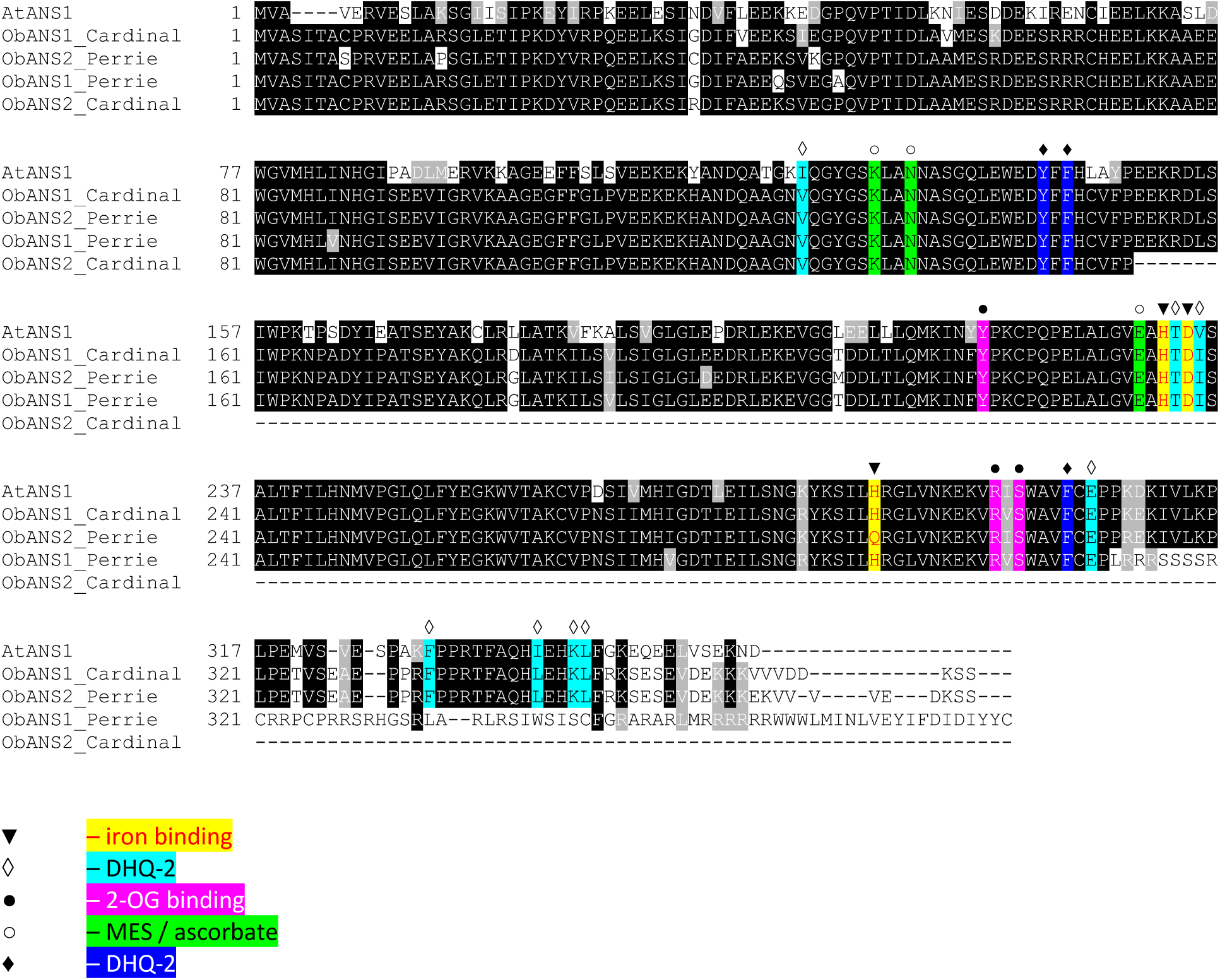
Multiple sequence alignment of anthocyanidin synthase proteins from sweet basil and Arabidopsis. Comparison between the amino acid sequences of basil ANS enzymes from ‘Perrie’ and ‘Cardinal’. Residues conserved in more than four sequence are black shaded and similar residues are gray shaded. Shapes and colors represent conserved residues based on the crystallographic structure of AtANS (Wilmouth et al., 2002). AtANS, *Arabidopsis thaliana*, accession number NP_194019.1.

### Models for ANS

To evaluate the effect of the mutations in the ANS enzymes of ‘Perrie’ and to estimate their contribution to the no anthocyanins phenotype, we have used a computational approach to predict their structure and functionality. First, we used Consurf to calculate the structural evolutionary conservation of each sequence residue in the target chain (Supplementary Fig. S4). The lowest score represents the most conserved position in a protein. The His-292 position in ANS proteins has a very low Consurf score of -1.3, indicating a highly evolutionary conserved amino acid. Some positions downstream to residue 311, where the frameshift of ANS1_Perrie occurred, are highly conserved from an evolutionary point of view which is often linked with functionally essential residues.

Next, we have constructed structure models for all four basil ANS enzymes. The Arabidopsis ANS crystal structure (1gp6; resolution of 1.75□), which shares ∼ 80% amino acid identities with basil ANSs, was used as a template for modeling calculated by RaptorX. First, we evaluated the influence of the H292Q substitution in ObANS2_Perrie on the active site and its possible effects on binding and activity. The electrostatic potential of the active site’s pocket is more negative in ObANS2_Perrie due to the substitution (Supplementary Fig. S5 AB), hence altering substrate and cofactors affinities. As observed in ANS crystallographic structure in the PDB database, the catalytic site populates an Fe^++^ ion as a cofactor, complexed with 2-oxoglutarate (2-OG) and the substrate t-dihydroquercetin (DHQ) to form the ANS:Fe(II):2OG:DHQ cluster (The natural substrate of ANS enzyme are leucoanthocyanidins, which are difficult to synthesize, and are unstable in a solution. Hence, the crystal structure of AtANS was resolved with DHQ). The Fe^++^ is coordinated with His-232, His-288, Asp-234, and a bidentate interaction with 2-OG in the 1pg6 structure, which are aligned with His-236, His-292 and Asp-238 respectively in the ‘Perrie’ and the ‘Cardinal’ variants (Supplementary Fig. S5 C). Three other coordination axes are held by two water molecules and a single succinate molecule. As indicated in the literature, the residues which mostly form coordination interactions with Fe^++^ ions are: Cys, Asp, Glu, His and Tyr, while Gln is less frequently participates in Fe^++^ coordination. The H292Q substitution might increase the affinity to other ions such as Ca^++^, K^++^, Na^++^, Mg^++^, Mn^++^ and Co^++^ (Zheng et al., 2008), which in turn would disturb the catalytic functionality and ANS:Fe(II):2OG:DHQ cluster formation.

To evaluate the frameshift effect in ObANS1_Perrie at the C-terminal tail, we used the structural model produced by RaptorX (Supplementary Fig. S6). The model predicts a rigid lobe composed of twisted alpha-helix, partially blocking the catalytic pocket (Supplementary Fig. S6). This kind of rigid structure might contain a hydrophobic core, which stabilizes the enzyme. The electrostatic potential of the frame-shifted C-terminal tail is different compared to the other variants and characterized by a more positive interface. Conservation analysis of the C-terminal raises a few conserved positions (Pro-311, Pro-320, His-341, Lys-345). Phe-334, Ile-338 and Leu-342 participate in DHQ-1 stabilization in the catalytic site and are absent due to frameshift predictably altering the enzyme’s catalytic activity.

### Complementation test in Arabidopsis

To validate the functionality of sweet basil ANS enzymes and the effect of the different sequences variation, we have introduced them into an Arabidopsis accession with a T-DNA insertion in its’ ANS gene. The plants of this Arabidopsis mutant do not accumulate anthocyanins at all in comparison to their Col-0 background (Bowerman *et al*., 2012). We have used all four *ObANS* genes and an ObANS2_Perrie with Q ⟶ H substitution in position 292 that presumably can fix the causative mutation and restore the enzyme’s activity. After floral dip *Agrobacterium*-mediated transformation, F1 plants were selected on Kanamycin-containing MS medium, grown on soil and seeds were harvested. F_2_ plants from four independent transformation events were germinated on Kanamycin-containing MS medium and the color of the plants was evaluated. Plants that acquired the *ObANS1_Cardinal* or the *ObANS2_Perrie_Q292H* genes showed an anthocyanin accumulating phenotype (Fig. 7). The anthocyanins were accumulated in the entire plants contrary to the Col-0 WT where they have been accumulated only in the center of the rosette. Contrary, plants acquired *ObANS2_Cardinal, ObANS1_Perrie* or *ObANS2_Perrie* showed a green phenotype similar to the phenotype of the Col-0 with T-DNA insertion at the *AtANS* gene.

**Figure 7.**
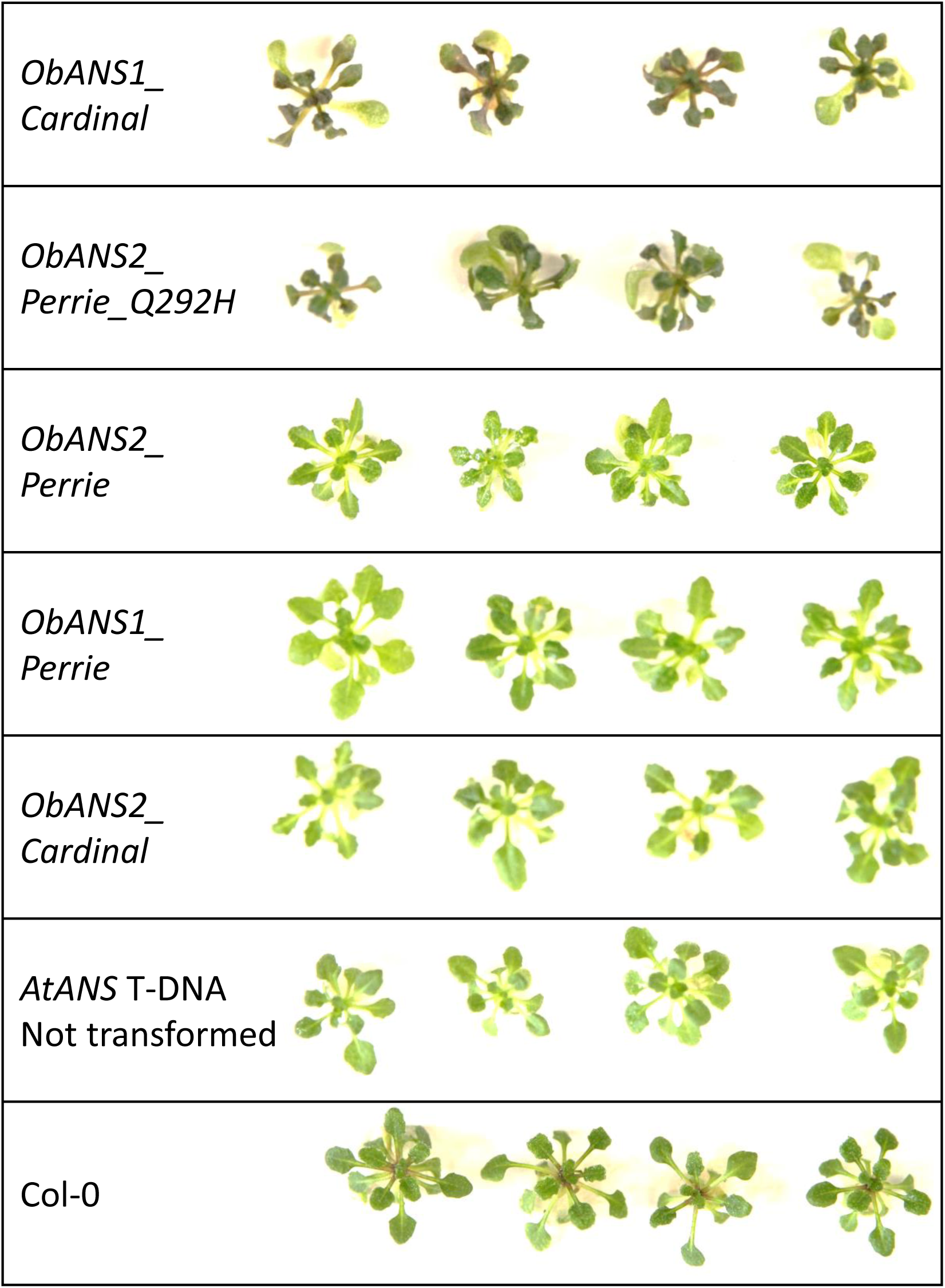
Complementation of the no anthocyanin phenotype in Arabidopsis by *ObANS* genes. The coding sequences of the four *ObANS* were extracted from cDNA, and cloned under the 35S promoter. The Q292H substitution in ObANS2_Perrie was performed using specific primers. The genes were transformed into Arabidopsis plants (Col-0 background) with a T-DNA insertion at the *AtANS* gene. The pictures depict F2 transgenes grown in MS medium with kanamycin at 20°C with continuous light. Each transgenic plant is representative of an independent transformation event.

## Discussion

### Color trait and anthocyanidin synthase

*Anthocyanidin synthase* encodes the first purple/blue/red color-committing enzyme in the anthocyanin biosynthetic pathway (Fig. 1). The function and role of ANS were demonstrated in multiple plants, including Arabidopsis (Appelhagen *et al*., 2011; Pelletier *et al*., 1997), apple (Szankowski *et al*., 2009), onion (Kim *et al*., 2004; Kim *et al*., 2005) grape (Gollop *et al*., 2001), and strawberry (Almeida *et al*., 2007). Multiple plants carry functional mutations in their ANS genes, causing non-colored phenotypes. For example, in pomegranates, it was shown that an insertion in the coding region of *PgANS* gene abolished anthocyanins accumulation in all plant parts (Ben-Simhon *et al*., 2015). In apple, it was shown that a viable ANS gene is critical for plant survival (Szankowski *et al*., 2009). Yet, in sweet basil, anthocyanins accumulation did not correlate with drought or salinity stress (Lazarević *et al*., 2021). Here we showed that two independent mutations in the homeologous genes of basil ANS are responsible for the all-green phenotype of commercial basil. Moreover, a non-functional allele of *ObANS2* was found in a purple-flower basil. The causing mutation is different from the non-functional mutation found in the green basil. Since the active ANS is dominant over the non-active ones, and there is a redundancy resulting from basil tetraploidy, it seems that a strong selection was applied towards green varieties, probably influenced by consumers’ demand.

While the mutations in the ANS genes clearly explain the green phenotype of ‘Perrie’ cultivar, they cannot explain the green leaves phenotype of the ‘Cardinal’ cultivar. Basils with purple leaves are commercially available and are very popular as ornamentals (Dudai and Belanger, 2016). They are not limited to a certain chemotype and, although more common in methyl-chavicol accumulating cultivars, purple eugenol and linalool basils exist (Liber *et al*., 2011; Maggio *et al*., 2016; Varga *et al*., 2017). Since ‘Cardinal’ and all progeny have green leaves, it seems that a different mechanism controls the color trait in the leaves in comparison to the flowers and stem. Phippen and Simon (2000) indicated a two dominant alleles inheritance mechanism for purple basil color. Since no segregant had purple leaves, another gene that alters anthocyanins accumulation in the leaves must exist. This gene might be an active repressor or defective inducer, but it seems that it cause green leaves in both cultivars. The expression of ANS genes is commonly regulated by MYB transcription factors in multiple plants (Xu *et al*., 2015; Zhang *et al*., 2014). “Non-purple” plants such as tomato carry a functional ANS gene as was demonstrated in tomato fruits, by exogenous expression of snapdragon transcription factors (Butelli *et al*., 2008), and by activation-tagged insertion lines (Mathews *et al*., 2003). We speculate that a similar mechanism is active in ‘Cardinal’ flowers and stems, and another different mechanism exists in the leaves of purple basil cultivars giving them their unique all-purple phenotype.

Arabidopsis mutants of the ANS gene have brown seeds (Abrahams *et al*., 2003; Abrahams *et al*., 2002). However, the seeds of ‘Perrie’ cultivar that has two non-functional ANS proteins are black as the seeds of ‘Cardinal’ cultivar and many other sweet basil accessions. That suggests, that in contrary to Arabidopsis, the dark color of basil seeds is not a result of anthocyanins accumulation. Several other phenolics are known to accumulate in seeds and have dark colors, including phlobaphenes accumulated in corn seeds and oxidized proanthocyanidins (Corso *et al*., 2020; Lepiniec *et al*., 2006). Yet, it seems that multiple metabolites contribute to the final color of the testa, as has been documented in *Brassica* (Yu, 2013). A detailed liquid-chromatography – mass-spectrometry analysis of basil seed coat may answer the question of which pigments contribute to the dark appearance of the testa in the lack of anthocyanins.

### Basil genomics and genetics

All color traits examined in this study showed disomic segregation with a dominant or incomplete dominance single gene inheritance model similar to the fusarium resistance trait (Gonda *et al*., 2022). That was also supported by the molecular findings of only a single functional ANS gene in ‘Cardinal’ cultivar. Moreover, the most proximate SNP to *ObANS1*, 7350_2,452,081, showed a disomic 1:2:1 distribution. On the contrary, the association mapping indicated a QTL on scaffold 120, where *ObANS2* is located. However, both *ObANS2_Perrie* and *ObANS2_Caridnal* are non-functional. Hence, the QTL observed at scaffold 120 can probably be explained by erroneous mapping of homeoloug reads to this scaffold instead of scaffold 7350. Support for this hypothesis can be found in the high level of heterozygosity detected in the ‘Cardinal’ parent, which was discussed by Gonda *et al*. (2022).

The large size of the QTL across the entire LG4A is in accordance with the small genetic distances found. It is currently unclear whether the large QTL is an artifact caused by genotyping errors, scaffolding errors, basil tetraploidy that causes a bioinformatic hurdle, or a real biology phenomenon. The parental lines might be too distant from each other, causing low recombination frequencies, resulting in small genetic distances and large QTLs. Large QTLs were observed in other corps, especially when located next to the centromere (Galpaz *et al*., 2018; Gao *et al*., 2019). Considering the very narrow QTL found for Fusarium wilt resistance in the same mapping population (Gonda *et al*., 2022), the later explanation should be considered.

### Expression of genes coding non-functional proteins

Another aspect of the current research is the high expression of genes encoding non-functional proteins. Both ‘Perrie’ and ‘Cardinal’ express inactive versions of the ANS gene. About 65% of the expressed *ObANS* RNA encoded mutant alleles. We have also observed an expression of mutated genes in the phenylpropenes biosynthesis pathway (Gonda et al., 2020). Phenotypically, polyploids can “allow” themselves to express mutated genes more than diploids since they have gene redundancy, as in the case of the flower color of ‘Cardinal’. Moreover, the selection against these mutated versions would be smaller because of this redundancy. However, this unnecessary expression and probable translation is futile and has energy costs. When dealing with biosynthetic pathways, such as anthocyanins biosynthesis, this cost is multiplied by the number of the genes involved. The fact that ‘Perrie’ was expressing the entire biosynthetic pathway in most of the cases of both homelogous caused even a greater futile expression effort. There is a long debate on the beneficiary of being a polyploid and its role in evolution and agriculture (Chen, 2010; Madlung, 2013). It has been suggested that polyploidy does not contribute to increased speciation and is an evolutionary dead-end (Mayrose *et al*., 2011; Wood *et al*., 2009). Yet, within domesticate plants, Salman-Minkov *et al*. (2016) showed that polyploidy promoted diversity and allowed wild polyploids to cope with agricultural niches and conditions.

Whatever will be the final answer in this debate, the degree of expression of genes encoding non-functional proteins and their possible energy cost should be considered. Whether it is beneficial for an organism to keep biosynthesis pathways “alive” at the expense of energy cost is another open question to be answered.

## Abbreviations

ANS: anthocyanidin synthase
LG: linkage group
QTL: quantitative trait loci

## Supplementary data

Table S1. Anthocyanins composition in ‘Cardinal’ cultivar vs. ‘Perrie’ cultivar.

Table S2. Contingency analysis of the phenotype within the purple segregants only.

Table S3. Scaffolds distribution to linkage groups.

Table S4. Chimeric scaffolds.

Table S5. Distribution of anthocyanins biosynthesis genes across sweet basil subgenomes.

Table S6. Expression of anthocyanins biosynthesis genes in ‘Perrie’ and ‘Cardinal’ flowers and leaves.

Table S7. Percentage identities of nucleotides of *ObANSs*.

Table S8. Percentage identities of amino acids of ObANS proteins and AtANS.

Fig. S1 Schematic illustration of the research population.

Fig. S2. Map charts of LG 1 and LG 4.

Fig. S3. Multiple sequence alignment of sweet basil ANS genes.

Fig. S4. Conserved residues analysis of ANS protein.

Fig. S5. A 3-D model structure of the active site of ObANS2_Perrie.

Fig. S6. A 3-D model structure for ObANS1_Perrie.

## Acknowledgments

The authors wish to thank lab and team members of the Unit of Aromatic and Medicinal Plants in Newe-Ya’ar Research Center in the past and present. Special thanks to Prof. Asaph Aharoni and Dr. Elad Oren for helpful discussion and brainstorming.

## Author contribution

RM, CA, AS, and MAA performed field and molecular biology experiments. DC and TKA performed and supervised field experiments. RDK and CA performed biochemistry analysis.

AF, OT and IG performed the bioinformatics analyses as well as the data curation. IG wrote the manuscript. IG and ND conceptualize the research and review the manuscript.

## Data availability

The raw data of the GBS is available through NCBI short reads archive (SRA) under project number: PRJNA836178.

