## Supplementary Fig. for "Two independent loss-of-function mutations in *anthocyanidin synthase* homeologous genes make sweet basil all green"

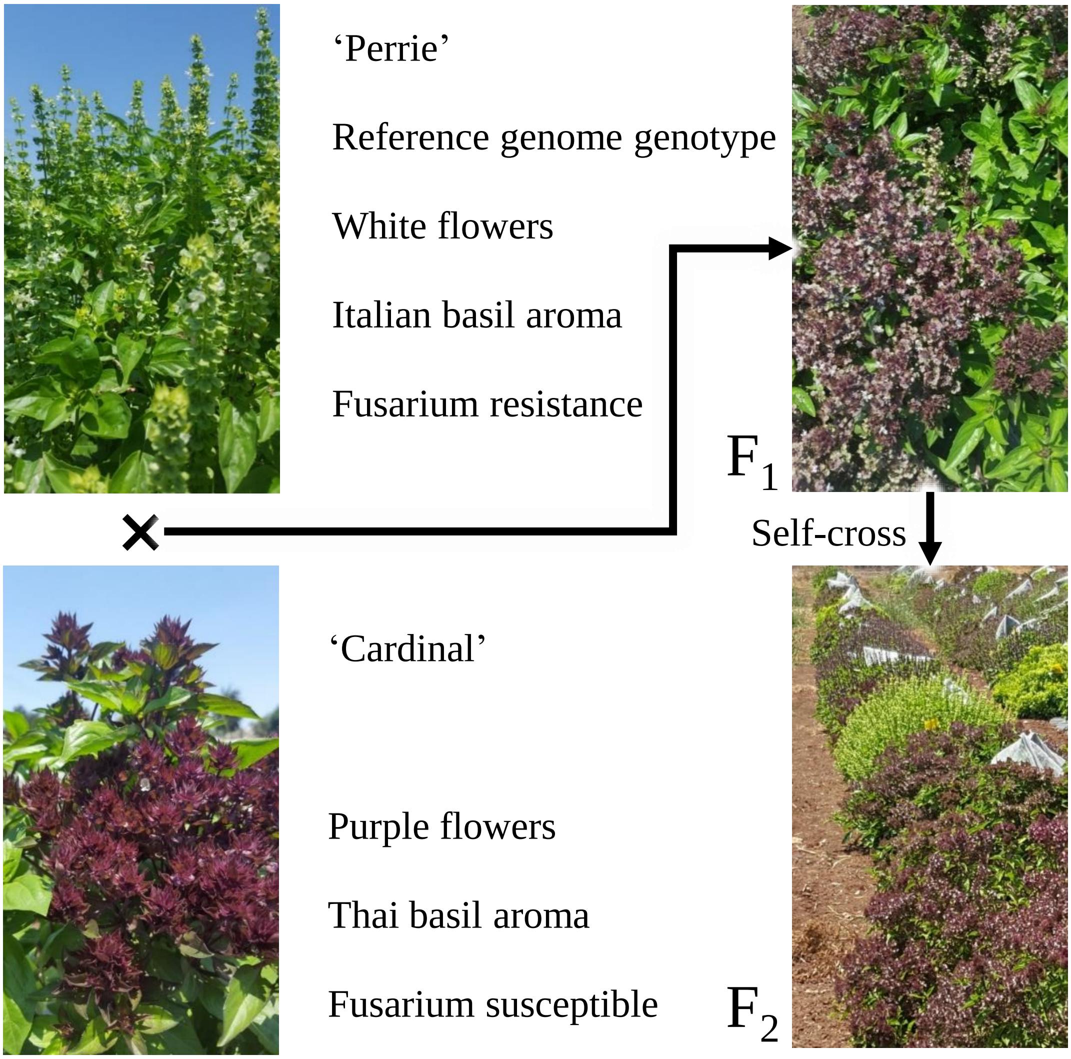


Figure S1. Schematic illustration of the studied population. Phenotypes of the parental lines and F1 generation plants are displayed.

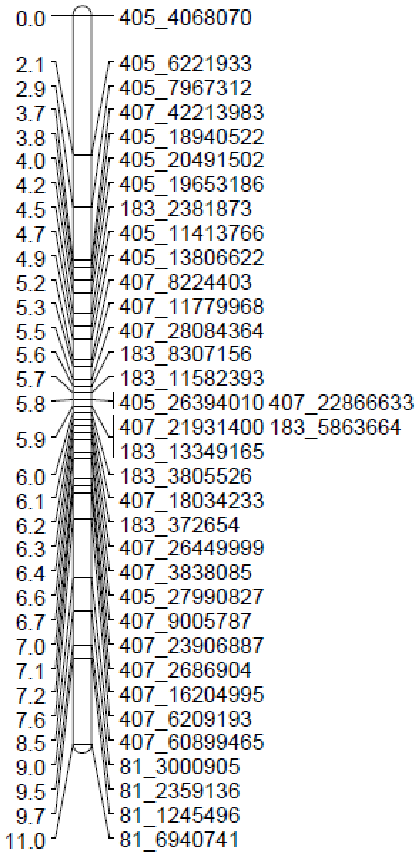


**Figure S2. Map charts of LG 1 and LG 4.** The maps generated display very short genetic distances as well as mixed order of the scaffolds.

```
ObANS1_Cardinal    1 ATGGTTGCTTCAATTACGGCATGCCCGCGGGTGGAGGAGCTGGCCCGAAGCGGGCTGGAGACGATCCCGAAAGACTACGT
ObANS1_Perrie      1 ATGGTTGCTTCAATTACGGCATGCCCTCGGGTGGAGGAGCTGGCCCGAAGCGGGCTGGAGACGATCCCCAAAGACTACGT
ObANS2_Cardinal    1 ATGGTTGCTTCAATTACGGCATGCCCGCGGGTGGAGGAGCTGGCCCGAAGCGGGCTGGAGACGATCCCGAAAGACTACGT
ObANS2_Perrie      1 ATGGTTGCTTCAATTACGGCAAGCCCGCGGGTGGAGGAGCTGGCCCCAAGCGGGCTGGAGACGATCCCGAAAGACTATGT

ObANS1_Cardinal   81 CCGGCCCCAAGAGGAGCTGAAGAGCATCGGCGACATATTCGTGGAAGAAAAGAGCATCGAGGGGCCGCAGGTTCCGACGA
ObANS1_Perrie     81 CCGGCCCCAAGAGGAGCTGAAGAGCATCGGCGACATATTCGCGGAAGAACAGAGCGTCGAGGGGGCGCAGGTTCCGACGA
ObANS2_Cardinal   81 CCGGCCTCAAGAGGAGCTGAAGAGCATCCGCGACATATTCGCCGAAGAAAAGAGCGTCGAGGGGCCGCAGGTTCCGACGA
ObANS2_Perrie     81 CCGGCCTCAAGAGGAGCTGAAGAGCATCTGCGACATATTCGCCGAAGAAAAGAGCGTCAAGGGGCCGCAGGTTCCGACGA

ObANS1_Cardinal  161 TCGATCTGGCGGTGATGGAGTCGAAAGACGAGGAGAGTCGGAGGAGGTGCCACGAGGAGCTGAAGAAGGCGGCGGAGGAG
ObANS1_Perrie    161 TCGATCTGGCGGCGATGGAGTCGAGAGACGAGGAGAGTCGGAGGAGGTGCCACGAGGAGCTGAAGAAGGCGGCGGAGGAG
ObANS2_Cardinal  161 TCGATCTGGCTGCGATGGAGTCGAGAGACGAGGAGAGTCGGAGGAGGTGCCACGAGGAGCTGAAGAAGGCGGCGGAGGAG
ObANS2_Perrie    161 TCGATCTGGCTGCGATGGAGTCGAGAGACGAGGAGAGTCGGAGGAGGTGCCACGAGGAGCTGAAGAAGGCGGCGGAGGAG

ObANS1_Cardinal  241 TGGGGGGTGATGCATCTGATCAACCATGGGATATCGGAGGAGGTGATCGGGCGCGTGAAGGCCGCCGGCGAGGGGTTTTT
ObANS1_Perrie    241 TGGGGGGTGATGCATCTGGTCAACCACGGGATATCGGAGGAGGTGATCGGGCGCGTGAAGGCCGCCGGCGAGGGGTTCTT
ObANS2_Cardinal  241 TGGGGGGTGATGCATCTGATAAACCACGGGATATCGGAGGAGGTGATCGGGCGCGTGAAGGCCGCCGGCGAGGGGTTCTT
ObANS2_Perrie    241 TGGGGGGTGATGCATCTGATAAACCACGGGATATCGGAGGAGGTGATCGGGCGCGTGAAGGCCGCCGGCGAGGGGTTCTT

ObANS1_Cardinal  321 CGGGCTGCCGGTGGAGGAGAAGGAGAAGCACGCCAACGATCAGGCCGCCGGAAACGTGCAGGGCTACGGCAGCAAGCTGG
ObANS1_Perrie    321 CGGGCTGCCCGTGGAGGAGAAGGAGAAGCACGCCAACGATCAGGCCGCCGGAAACGTGCAGGGCTACGGCAGCAAGCTGG
ObANS2_Cardinal  321 TGGGCTGCCCGTGGAGGAGAAGGAGAAGCACGCCAACGATCAGGCCGCCGGAAACGTGCAGGGCTACGGCAGCAAGCTGG
ObANS2_Perrie    321 TGGGCTGCCCGTGGAGGAGAAGGAGAAGCACGCCAACGATCAGGCCGCCGGAAACGTGCAGGGCTACGGCAGCAAGCTGG

ObANS1_Cardinal  401 CGAATAACGCTAGCGGGCAGCTGGAGTGGGAGGATTATTTCTTTCACTGTGTTTTTCCGGAGGAGAAGAGGGACTTGTCT
ObANS1_Perrie    401 CGAATAACGCTAGCGGGCAGCTGGAGTGGGAGGATTATTTCTTTCACTGTGTTTTTCCGGAGGAAAAGAGGGACTTGTCC
ObANS2_Cardinal  401 CGAATAACGCTAGCGGGCAGCTCGAGTGGGAGGATTATTTCTTTCACTGTGTTTTCCCGTAGGAGAAGAGAGACTTGTCT
ObANS2_Perrie    401 CGAATAACGCTAGCGGGCAGCTGGAGTGGGAGGATTATTTCTTTCACTGTGTTTTCCCGGAGGAGAAGAGGGACTTGTCT

ObANS1_Cardinal  481 ATTTGGCCCAAGAATCCGGCTGATTATATACCGGCAACAAGCGAGTATGCAAAGCAACTAAGAGACCTAGCAACCAAGAT
ObANS1_Perrie    481 ATTTGGCCCAAGAATCCGGCTGATTATATACCGGCAACAAGCGAGTATGCAAAGCAACTAAGAGGCCTAGCAACCAAGAT
ObANS2_Cardinal  481 ATTTGGCCCAAGAATCCGGCTGATTATATAACGGCAACAAGCGAGTATGCAAAGCAACTAAGAGGCCTAGCAACCAAGAT
ObANS2_Perrie    481 ATTTGGCCCAAGAATCCGGCTGATTATATACCGGCAACAAGCGAGTATGCAAAGCAACTAAGAGGCCTAGCAACCAAGAT

ObANS1_Cardinal  561 CCTCTCAGTCCTCTCAATCGGCCTTGGATTAGAAGAAGACCGCCTAGAAAAAGAAGTCGGCGGAACGGACGATCTTACCC
ObANS1_Perrie    561 CCTCTCAGTCCTCTCAATCGGCCTTGGATTAGAAGAAGACCGCCTAGAAAAAGAAGTCGGCGGAACGGACGATCTTACCC
ObANS2_Cardinal  561 CCTCTCGATCCTCGCAATCGGCCTTGGACTAGACGAAGACCGCCTAGAAAAAGAAGTCGGCGGAATGGACGATCTTACTC
ObANS2_Perrie    561 CCTCTCGATCCTCTCAATCGGCCTTGGACTAGACGAAGACCGCCTAGAAAAAGAAGTCGGCGGAATGGACGATCTTACCC

ObANS1_Cardinal  641 TCCAGATGAAGATAAACTTCTACCCGAAATGCCCTCAGCCGGAGCTGGCCCTTGGCGTGGAGGCCCACACCGACATCAGC
ObANS1_Perrie    641 TCCAGATGAAGATAAACTTCTACCCGAAATGCCCTCAGCCGGAGCTGGCCCTCGGCGTGGAGGCCCACACCGACATCAGC
ObANS2_Cardinal  641 TCCAGATGAAGATAAACTTCTACCCGAAATGCCCTCAGCCGGAGCTGGCCCTCGGCGTGGAGGCCCACACCGACATCAGC
ObANS2_Perrie    641 TCCAGATGAAGATAAACTTCTACCCGAAATGCCCTCAGCCGGAGCTGGCCCTCGGCGTGGAGGCCCACACCGACATCAGC

ObANS1_Cardinal  721 GCCCTCACCTTCATCCTCCACAACATGGTCCCCGGCCTGCAGCTCTTCTACGAGGGCAAATGGGTCACTGCCAAATGTGT
ObANS1_Perrie    721 GCCCTCACCTTCATCCTCCACAACATGGTCCCCGGCCTGCAGCTCTTCTACGAGGGCAAATGGGTCACTGCCAAATGCGT
ObANS2_Cardinal  721 GCCCTCACCTTCATCCTCCACAACATGGTCCCCGGCCTGCAGCTCTTCTACGAAGGCAAATGGGTCACTGCCAAATGTGT
ObANS2_Perrie    721 GCCCTCACCTTCATCCTCCACAACATGGTCCCCGGCCTGCAGCTCTTCTACGAGGGCAAATGGGTCACTGCCAAATGTGT

ObANS1_Cardinal  801 GCCCAACTCCATCATCATGCACATCGGCGACACCATCGAGATTCTCAGCAACGGCAGATACAAAAGCATTCTGCACCGCG
ObANS1_Perrie    801 GCCCAACTCCATCATCATGCACGTCGGCGACACCATCGAGATTCTCAGCAACGGCAGATACAAAAGCATTCTGCACCGCG
ObANS2_Cardinal  801 GCCCAACTCCATCATCATGCACATCGGCGACACCATCGAGATTCTCAGTAACGGCAAATACAAAAGCATTCTGCACCGCG
ObANS2_Perrie    801 GCCCAACTCCATCATCATGCACATCGGCGACACCATCGAGATTCTCAGTAACGGCAAATACAAAAGCATTCTGCAGCGCG

ObANS1_Cardinal  881 GCCTCGTCAACAAGGAGAAGGTGAGGGTTTCTTGGGCGGTTTTCTGCGAGCCGCCTAAGGAGAAGATTGTGCTCAAGCCG
ObANS1_Perrie    881 GCCTCGTCAACAAGGAGAAGGTGAGGGTTTCTTGGGCGGTTTTCTGCGAGCC-CCTAAGGAGAAGATCGTCCTCAAGCCG
ObANS2_Cardinal  881 GCCTCGTCAACAAGGAGAAGGTGAGGATTTCTTGGGCGGTTTTCTGCGAGCCGCCTAAGGAGAAGATCTTCCTCAAGCCG
ObANS2_Perrie    881 GCCTCGTCAACAAGGAGAAGGTGAGGATTTCTTGGGCGGTTTTCTGCGAGCCGCCTAGGGAGAAGATCGTCCTCAAGCCG

ObANS1_Cardinal  961 CTGCCGGAGACCGTGTCCGAGGCGGAGCCGCCGCGGTTCCCGCCGCGCACGTTTGCGCAGCATCTGGAGCATAAGCTGTT
ObANS1_Perrie    960 CTGCCGGAGACCGTGTCCGAGGCGGAGCCGCCACGGTTCCCGCCTCGCACGTTTGCGCAGCATCTGGAGCATAAGCTGTT
ObANS2_Cardinal   961 CTGCCCGAGACCGTGTCCGAGGCGGAGCCGCCGCGGTTCCCTCCGCGCACGTTTGCGCAGCATCTGGAGCATAAGCTGTT
ObANS2_Perrie    961 CTGCCGGAGACCGTGTCTGAGGCGGAGCCGCCGCGGTTCCCTCCGCGCACGTTTGCACAGCATCTGGAGCATAAGCTGTT

ObANS1_Cardinal 1041 TAGGAAGAGCGAGAGCGAGGTTGATGAGAAGAAGAAG---------GTGGTCGTTGATGATAAATCTAGTTGA-------
ObANS1_Perrie   1040 TCGGAAGAGCGAGAGCGAGGTTGATGAGAAGAAGAAGGAGAAGGTGGTGGTGGTTGATGATAAATCTAGTTGAATATATA
ObANS2_Cardinal 1041 TAGGAAGAGCGAGAGCGAGGTTGATGAGAAGAAGAAGGAGAAGGTGGTGGGGGTTGG--ATGA-----------------
ObANS2_Perrie   1041 TAGGAAGAGCGAGAGCGAGGTTGATGAGAAGAAGAAGGAGAAGGTGGTGGTGGTTGAGGATAAATCTAGTTGA-------

ObANS1_Cardinal      ---------------------------
ObANS1_Perrie   1120 TTTGATATTGATATTTACTATTGTTGA
ObANS2_Cardinal      ---------------------------
ObANS2_Perrie        ---------------------------
```

Figure S3. Multiple sequence alignment of sweet basil ANS genes. Sequence were derived from cDNA clones. Alignment was performed with ClustalOmega (Madeira et al., 2019).

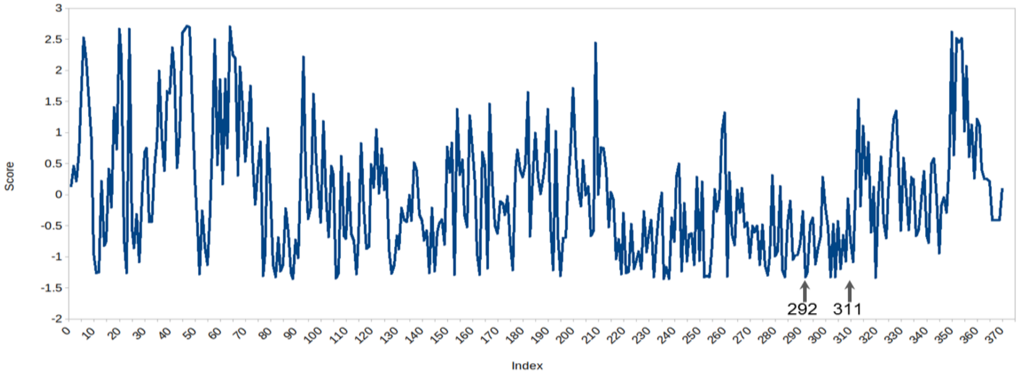


Figure S4. Conserved residues analysis of ANS protein. Evolutionary conserved amino acid residues were estimated using Consurf.

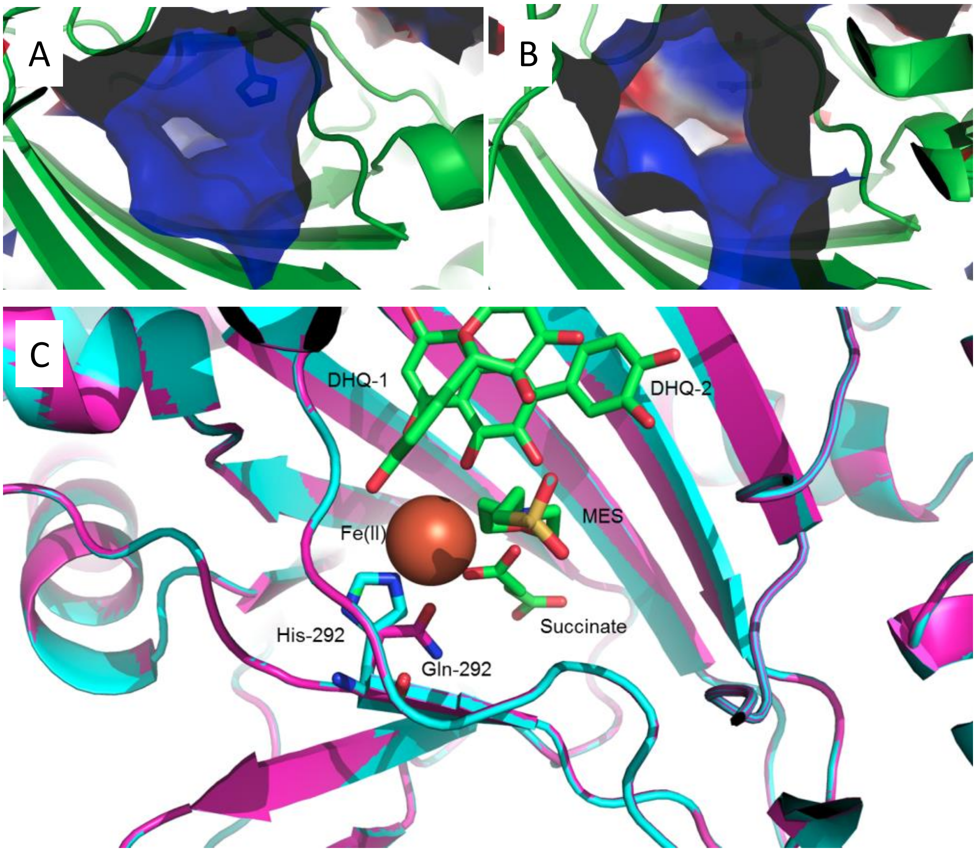


Figure S5. A 3-D model structur eof the active site's of ObANS2_Perrie. The surface electrostatic potential of the active site's pocket of ObANS1_Cardinal (A) and ObANS2_Perrie (B). Negative potential is colored in red and positive potential is colored in blue. C. Structure models produced by RaptorX of the active site of ObANS2_Perrie (Magenta) and ObANS1_Cardinal (Cyan). The H292Q substitution is represented in sticks. The crystal structure 1pg6 was superimposed on the ANS models, including DHQ-1, 2OG, Fe(II) and 2OG.

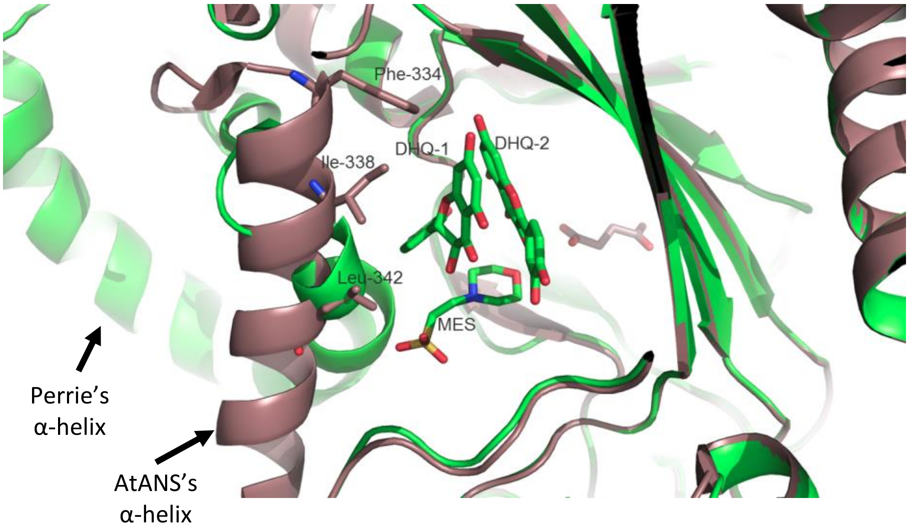


Figure S6. A3-D model structure for ObANS1_Perrie. Focus on the catalytic site of AtANS (Brown) where DHQ-1 is stabilized by hydrophobic interactions of Phe-334, Ile-338 and Leu342. ObANS1_Perrie variant (green) presents a twisted alpha helix absent these residues.
